# hnRNP R is a regulator of stress granule formation

**DOI:** 10.1101/2022.06.07.495171

**Authors:** Changhe Ji

## Abstract

hnRNP R (Heterogeneous nuclear ribonucleoproteins R) is one member of RNA-binding proteins (RBPs) in hnRNP family, it plays important role in nucleic acid metabolism including alternative splicing, mRNA stabilization, transcriptional and translational regulation, and mRNA transport. Mostly hnRNP R is localized in nuclear and extensively binds with RNA and chromatin, but still some part of it exports to cytosol. We found that long and short isoform hnRNP R can located in stress granules, depletion hnRNP R can facilitate the stress granule formation and accelerate mRNA in the stress granules. Meanwhile we found that depletion hnRNP R can alter the stress granule phenotype and impair stress granule resolving. Additionally, we also found that depletion hnRNP R also induce TDP43 granules formation. Furthermore, depletion hnRNP R can impair TDP43 binding with ribosome. Altogether, hnRNP R plays important role in the stress granule formation.

## INTRODUCTION

Stress granules (SGs) are nonmembrane assemblies formed in cells in response to stress conditions, such as chemical, heat shock, and virus invasion etc. SGs mainly contain untranslated mRNA and a variety of proteins [1] [2]. Stress granules are thought to influence mRNA function, localization, and to affect signaling pathways [3] [4] [5]. Normally, stress granule formation is a dynamic, reversible process. However, pathological mutations in proteins that either increase the formation, or decrease the clearance of stress granules, can lead to abnormal accumulation of aggregates that share components with stress granules [6] [8] [9]. Abnormal accumulation of stress granule-like aggregates is associated with neurodegenerative disease [10],such as ALS, FTD, HD and AD [11-13]. The molecular interactions and mechanisms that regulate stress granule assembly and how these may become altered in disease remain inclusive. Anyhow, the mechanism behind stress granules whether have connection with pTDP-43 aggregation still not clear and need to be investigated further.

Heterogeneous nuclear ribonucleoproteins (hnRNPs) are a family of functionally diverse RNA bindings proteins (RBPs) [14]. Originally named alphabetically from A1 to U, they range from 34 to 120 kDa [15]. Different with most of other hnRNPs, hnRNP R has two isoforms. Both isoforms localize in the nuclear, but some still can exist in the cytosol and motoneuron axons and growth cones[16] [17] [18]. hnRNP R’s function also accompany binding with other RNAs, such as 7SK, both were found can coregulate the axonal transcriptome of motoneurons[19]. hnRNP R was also found binding with SMN in 7SK dependent manner[20], this indicates SMN’s dysfunction also affect hnRNP R normal functions. Also, loss of full-length hnRNP R isoform impairs DNA damage response[21], which is not surprising, because mostly hnRNP R binds with chromatin and nocent RNA. Besides hnRNP R plays important role in normal cell physiological function, hnRNP R also associates with motoneuron degenerative disease, FUS mutation caused FTLD has been found that hnRNP R can form and accumulate in pathological inclusions with FUS [18]. Furthermore, one study found that de novo truncating and missense variants in HNRNPR in four unrelated individuals presenting with overlapping neurodevelopmental phenotypes and dysmorphic features[22].

Above mentioned is the summary of the stress granule and hnRNP R’s function. But one important question is what is hnRNP R’s role in stress granule formation still not answer yet. Here, we found that depletion hnRNP R can facilitate the stress granule formation and alter stress granule phenotype, meanwhile impair stress granule resolving. Altogether, those results give us an idea that hnRNP R, to some exitance can inhibit the TDP43 granule formation. Specifically, a better understanding of TDP-43 aggregates formation will surely shed light on novel therapeutics that have the potential to most ALS cases.

## RESULTS

### 1. MePCE, but not LARP7, HEXIM1, CDK9, Cyclin T1 localized in stress granule

7SK recruits HEXIM1 and p-TEFB to form 7SK/p-TEFB complex which can inhibit p-TEFB transcription activity in the nuclear[23], besides this complex, 7SK can bind with hnRNP proteins to form another separate complex, which is called 7SK/hnRNP complex[24]. The dynamics balance of these two complexes plays important role in RNA and protein homeostasis

At the beginning, we checked 7SK/p-TEFB complex associated protein whether goes to stress granules or not. Very interestingly, we found that MePCE, but not LARP7, HEXIM1, CDK9, cyclin T1 located in the stress granule (figure 1, supplementary figure 1). MePCE contains a methyltransferase domain and is responsible for adding a unique γ-monomethyl phosphate cap structure onto the 5′-end of 7SK[25]. LARP7 is another stable component of the 7SK snRNP[26], MePCE and LARP7 forms the core of 7SK snRNP [27] [28] [29] [30] [31]. This different role between MePCE and LARP7 in stress granules, indicates that MePCE can release from 7SK complex without disturb the stability of 7SK. This result is consistent with previous finding that MePCE is one competent of stress granules but not LARP7[32]. Although HEXIM1 is one competent of 7SK/P-TEFb complex [33] [34], but it does not located in stress granules meanwhile we also found that CDK9 and cyclin T1 do not go to stress granules. Taken together, we found that the muti-roles of 7SK complex associated proteins in the cytoplasm besides nuclear.

**Figure 1.**
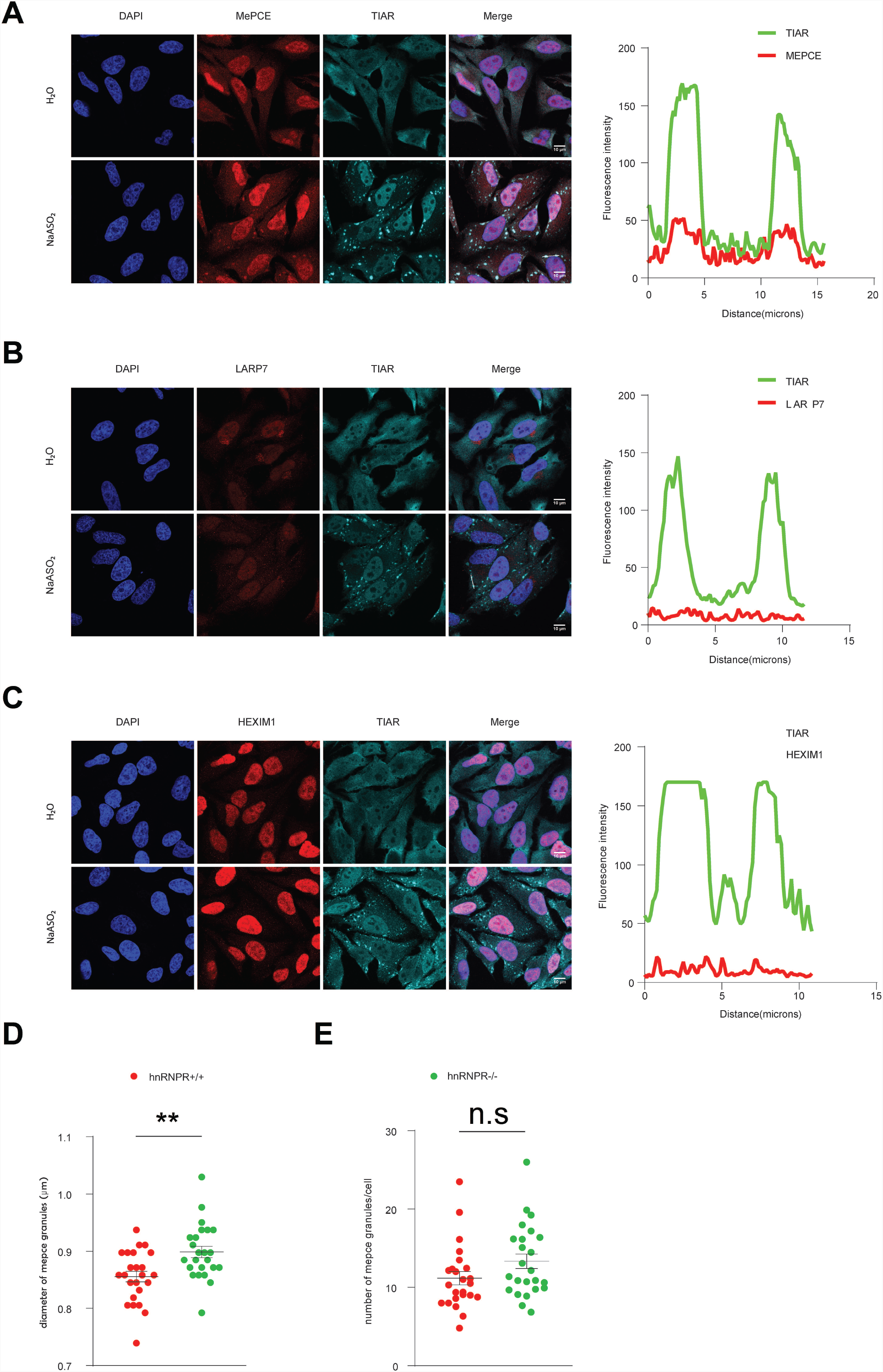
MEPCE, but not LARP7, HEXIM1, CDK9, Cyclin T1 localized in stress granule. A Immunostaining with antibodies against MePCE and TIAR. Scale bar: 10 μm. B Immunostaining with antibodies against LARP7 and TIAR. Scale bar: 10 μm C Immunostaining with antibodies against HEXIM1 and TIAR. Scale bar: 10 μm D Quantification of the diameter of MePCE granules in per *HNRNPR*+/+ and *HNRNPR*-/- cells Data are mean with standard deviation (SD); \*\**P* ≤ 0.01; unpaired two-tailed *t*- test (n = 3 biological replicates). E Quantification of the number of MePCE granules in per *HNRNPR*+/+ and *HNRNPR*-/- cells Data are mean with standard deviation (SD); \*\**P* ≤ 0.01; unpaired two-tailed *t*-test (n = 3 biological replicates).

### 2. hnRNP R localized in stress granule

After finding 7SK/P-TEFb complex does not go to stress granules, next we move to 7SK/hnRNP complex associated proteins whether go to stress granules or not. hnRNP R is one of deeply validated hnRNP proteins which is belong to 7SK/hnRNP complex by many groups [35] [36] [37] [38] [39] [29] [27] [40] [41] [42] [43]. We treated the Hela and HEK293T cells with sodium arsenate then used an antibody which target c terminal of hnRNP R and stain with the stress granule marker TIAR [44] [45] [46], hnRNP R can clearly colocalize with TIAR (figure 2), we used another antibody which target N terminal of long isoform of hnRNP R, and found that hnRNP R long isoform also goes to the stress granules(supplementary figure 2). Taken together those results confirm that hnRNP R long isoform localized in stress granule.

**Figure 2.**
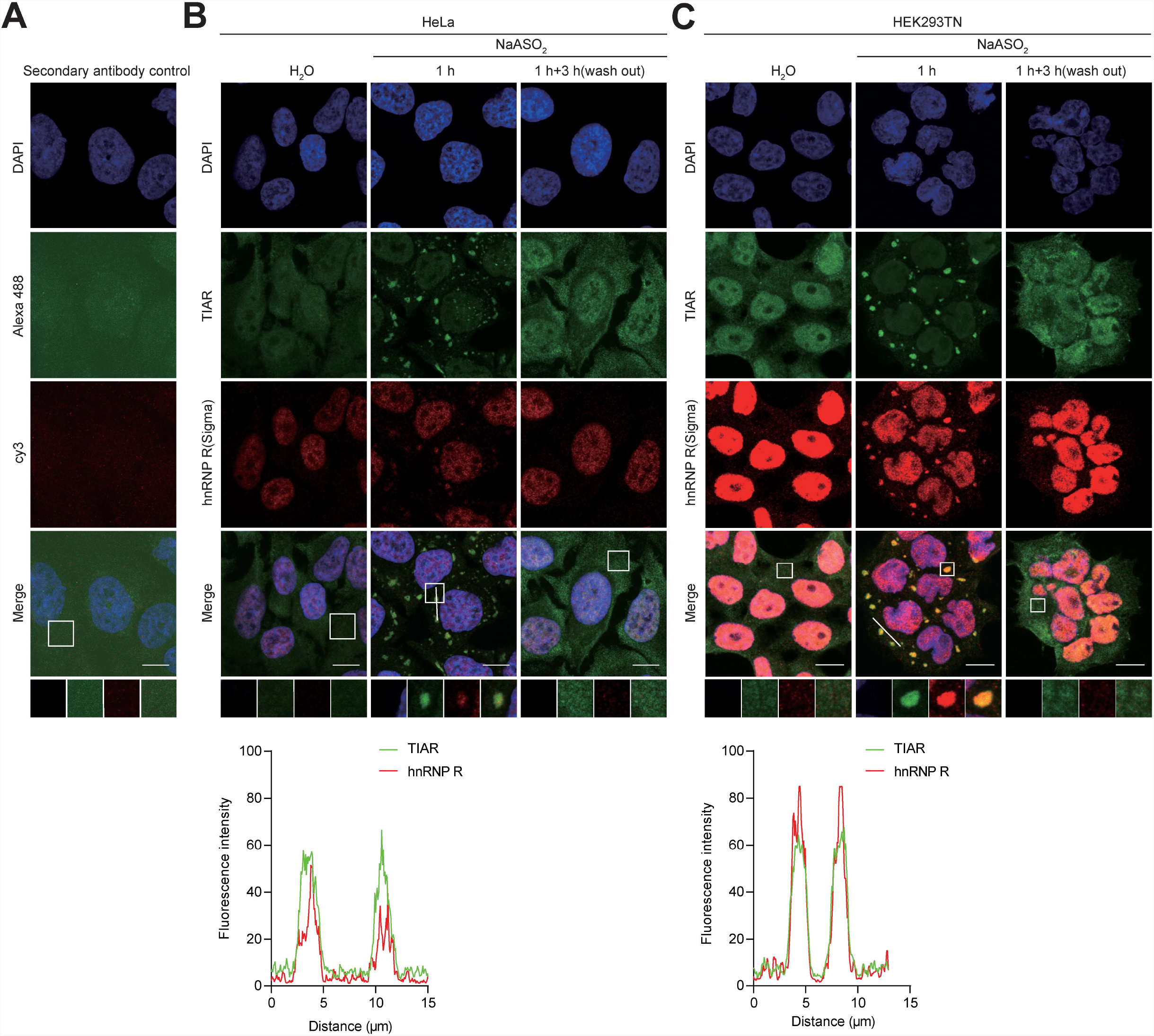
hnRNP R localized in stress granule. A Immunostaining with secondary antibodies control. Scale bar: 10 μm.. B Immunostaining with antibodies against TIAR and hnRNP R in HeLa cells with or without sodium arsenite treatment and wash out sodium arsenite. Scale bar: 10 μm C Immunostaining with antibodies against TIAR and hnRNP R in HEK293TN cells with or without sodium arsenite treatment and wash out sodium arsenite. Scale bar: 10 μm.

### 3. hnRNP R long and short isoform localized in stress granule

Using commercial antibody, we confirmed long hnRNP R located in the stress granules, but we still cannot confirm that the short isoform went to the stress granules or not.

To distinguish different isoforms of hnRNP R, a GFP tagged hnNRP R was added in the c terminal of long and short hnRNP R, and then cloned into pcDNA 3.0 vector, HeLa cells was transfected and harvest cells after 48 hours, then load the same amount of total protein for western. As shown in supplementary figure 3B and 3C, GFP, hnRNP R_long_GFP, hnRNP R_short_GFP was expressed in correct size. We also found that GFP tagged long isoform of hnRNP R was expressed less than short isoform. Additionally we used a nano-trap GFP agarose beads to pull down GFP or GFP tagged hnRNP R, as shown in supplementary figure 3D, GFP or GFP tagged hnRNP R was successfully pull down by western, meanwhile, we found 7SK and malat1 ncRNA was both highly binding with hnRNP R by doing RIP-qPCR(supplementary figure 3E), this result is consistent with previous finding [37].Very interestingly, long isoform hnRNP R protein was less pulling down, long isoform hnRNP R can pull down more 7SK than short isoform hnRNP R, this means that long and short isoform hnRNP R belong to different 7SK complex.

After validation the plasmids, next we transfected GFP tagged long and short isoform hnRNP R plasmids in HeLa and HEK293T cells and treat the cells with sodium arsenate, then stain GFP and TIAR, we found that both long and short isoform hnRNP R went to the stress granules (Figure 3).

**Figure 3.**
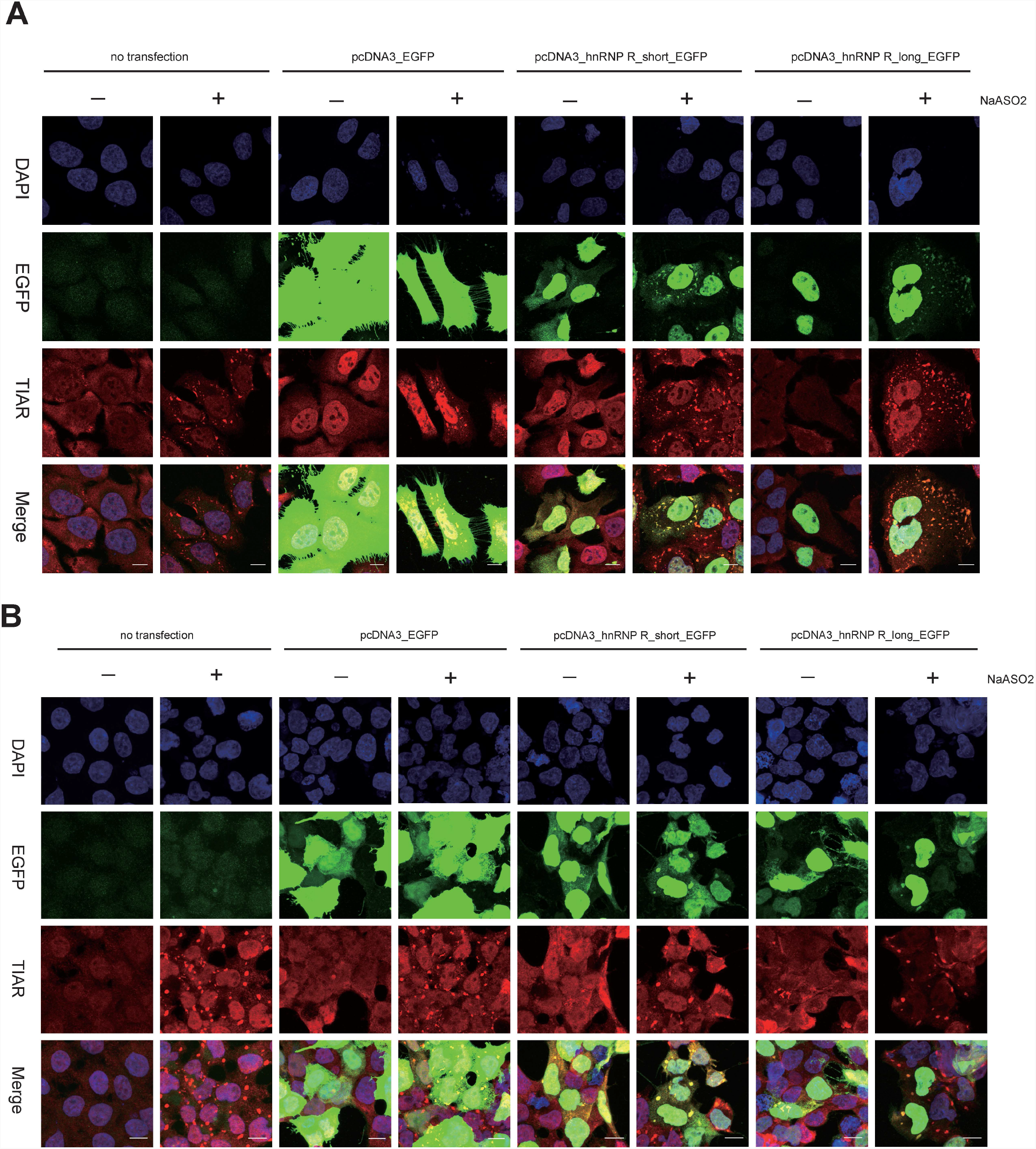
hnRNP R long and short isoform localized in stress granule. A HeLa cells without transfection or transfected with pcDNA3_EGFP, pcDNA3_hnRNP R_short_EGFP,, pcDNA3_hnRNP R_long_EGFP plasmid, then treat with water or 0.5mM sodium arsenite, then Immunostaining with antibodies against EGFP and TIAR, Scale bar: 10 μm.. B HEK293TN cells without transfection or transfected with pcDNA3_EGFP, pcDNA3_hnRNP R_short_EGFP,, pcDNA3_hnRNP R_long_EGFP plasmid, then treat with water or 0.5mM sodium arsenite, then Immunostaining with antibodies against EGFP and TIAR, Scale bar: 10 μm.

### 4. depleting hnRNP R stimulates stress granule formation

Although previous study has found hnRNP R localized in the stress granules[22, 47]. No study systematic investigated the hnRNP R’ s function in the stress granules formation. Based on this question, we depleted endogenous hnRNP R by using lentivirus derived shRNA, meanwhile control using GFP virus and we also validated the knockdown efficiency by qPCR, western blot and staining (figure 4 and supplementary figure 2),all get around 90% knockdown efficacy, after that, we treat the Hela cells which are transduced with GFP or sh*HNRNPR* knockdown virus with sodium arsenate, as control we use water, very interestingly, we found that, even without sodium arsenate treatment, the cells can also form stress granules after knockdown hnRNP R (figure 4), this can be explained that without hnRNP R, the cells undergo a lot of metabolic disorders, but the mechanism still need to be investigated, one recent study found that knock down RNF4 or SUMO2/3 also can stimulate stress granules formation [47]. This is not surprised, any factor which connected with stress granules is depleted will make the cells stress.

**Figure 4.**
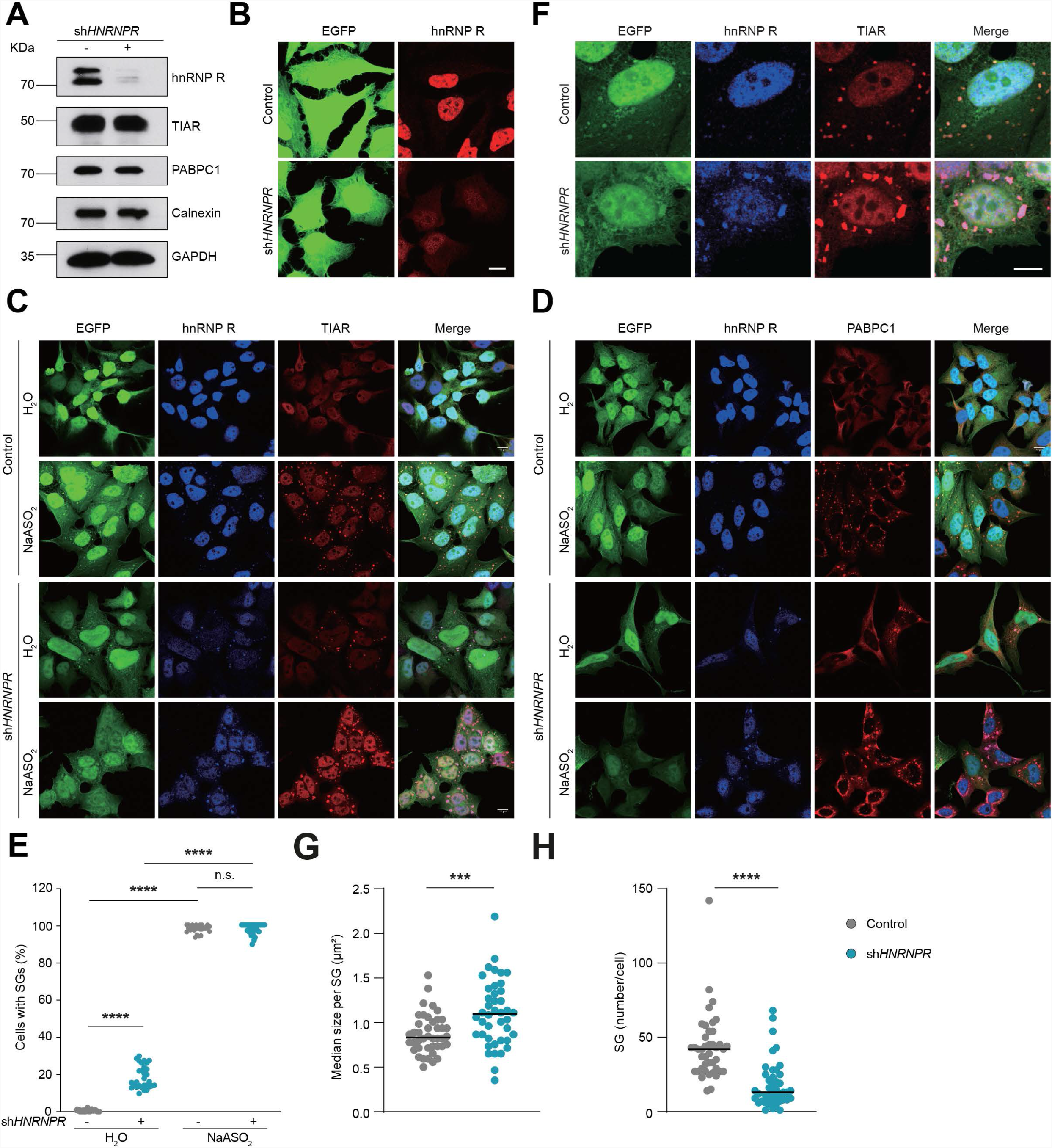
depleting hnRNP R stimulates stress granule formation. A Western blot analysis of hnRNP R, TIAR, PABPC1, Calnexin and GAPDH in control and sh*HNRNPR* cell lysate. B Immunostaining with antibodies against GFP and hnRNP R in control and sh*HNRNPR* cell. Scale bar: 10 μm. C Immunostaining with antibodies against GFP, TIAR and hnRNP R in control and sh*HNRNPR* cell without or with sodium arsenite treatment. Scale bar: 10 μm. D Immunostaining with antibodies against GFP, PABPC1 and hnRNP R in control and sh*HNRNPR* cell without or with sodium arsenite treatment. Scale bar: 10 μm. E Quantification of the percentage of stress granule positive cells in control and sh*HNRNPR* cell without or with sodium arsenite treatment. F Immunostaining with antibodies against GFP, TIAR and hnRNP R in control and sh*HNRNPR* cell with sodium arsenite treatment. Scale bar: 10 μm. G Quantification of the median size per stress granules in per control and sh*HNRNPR* cell. Data are mean with standard deviation (SD); \*\*\**P* ≤ 0.001; unpaired two-tailed *t*- test (n = 3 biological replicates). H Quantification of the number of stress granules in per control and sh*HNRNPR* cell. Data are mean with standard deviation (SD); \*\*\*\**P* ≤ 0.0001; unpaired two-tailed *t*-test (n = 3 biological replicates).

### 5. depleting hnRNP R altered RNA granule and stress granules phenotype

CRISPR/cas9 technology has become the most popular genome editing tool since it was first used in 2012 [48], during 9 years, hundreds of new technology surrounding CRISPR/cas9 has comes out, including Prime editing [49], this technology overcome lots of defects which were previous used for CRISPR/cas9, such as no need donor DNA, only cut single strand of DNA which can dramatically reduce off target effect. people use prime editing to knock in a small fragment at any site as you wish theoretically without deleting large fragment, If completely knockout hnRNP R need cut nearly 3kb DNA region without promising the accuracy. Here we knock in a AGTGA five nuclear acids and introduce a stop codon at exon 4 of long hnRNP R and exon 3 of short hnRNP R. The cutting efficiency is around 50%, including 40% heterozygous colonies and 10% homozygous colonies, our knockout efficiency is higher than the lecture reported [49], this is can be explained by different genes has different cutting efficiency.

Next step, we checked RNA granule and stress granules phenotype after depletion hnRNP R we found that RNA granule and stress granules are becomes bigger and less in *HNRNPR* knockdown or knockout cells (Figure 4, 5, 6, 7 and supplementary figure 4).

**Figure 5.**
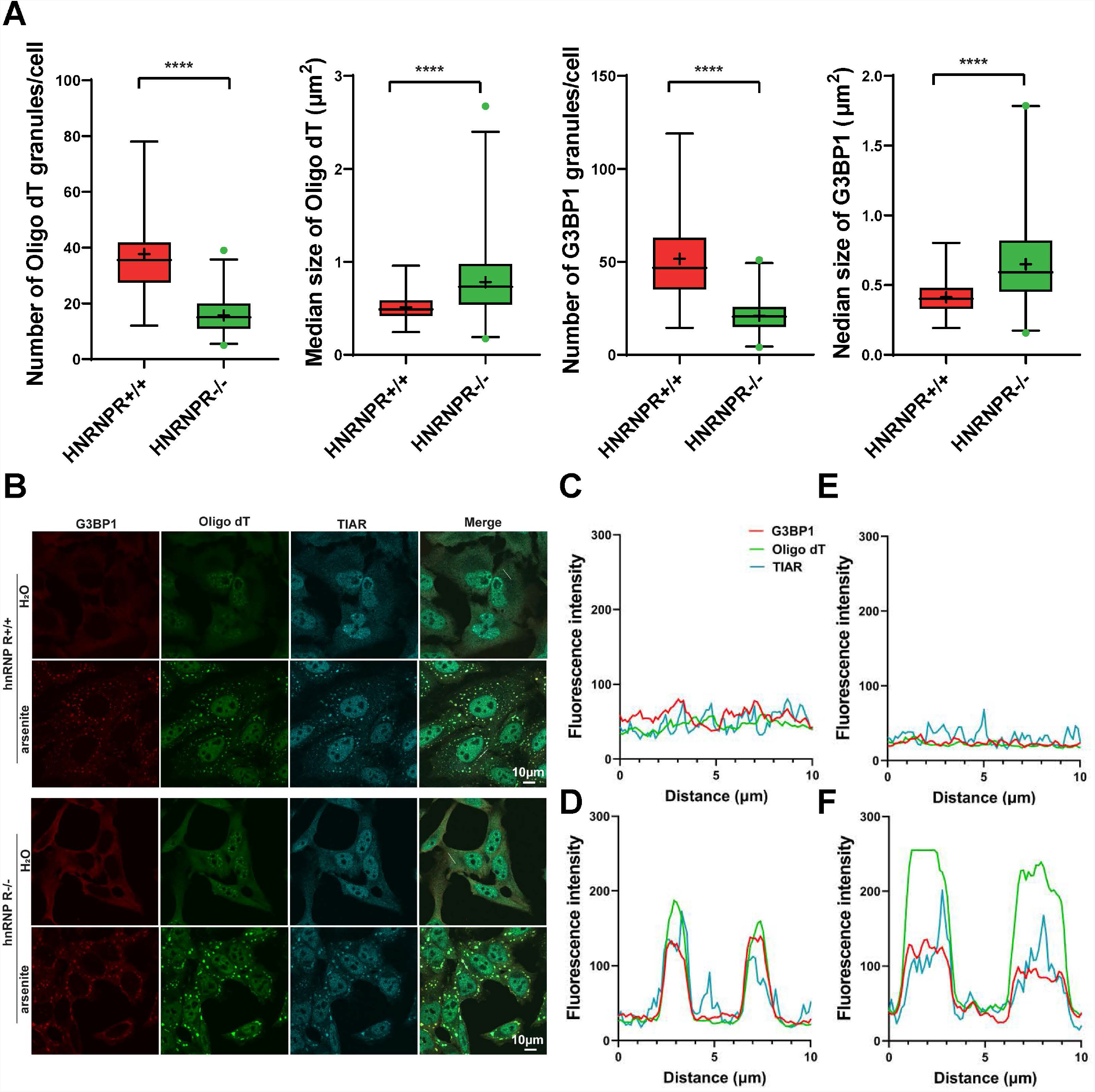
depleting hnRNP R facilitates RNA granule formation. A Quantification of the number and the size of RNA granules and stress granules in per in *HNRNPR* +/+ and -/- cells. Data are mean with standard deviation (SD); \*\*\*\**P* ≤ 0.0001; unpaired two-tailed *t*-test (n = 3 biological replicates). B Poly(A) FISH and Immunostaining with antibodies against TIAR and G3BP1 in *HNRNPR* +/+ and -/- cells with or without sodium arsenite treatment. Scale bar: 10 μm. C, D TIAR, G3BP1 and oligo-dT fluorescence intensity curve in *HNRNPR* +/+ cells without or with sodium arsenite treatment. Scale bar: 10 μm. E, F TIAR, G3BP1 and oligo-dT fluorescence intensity curve in *HNRNPR* -/- cells without or with sodium arsenite treatment. Scale bar: 10 μm.

**Figure 6.**
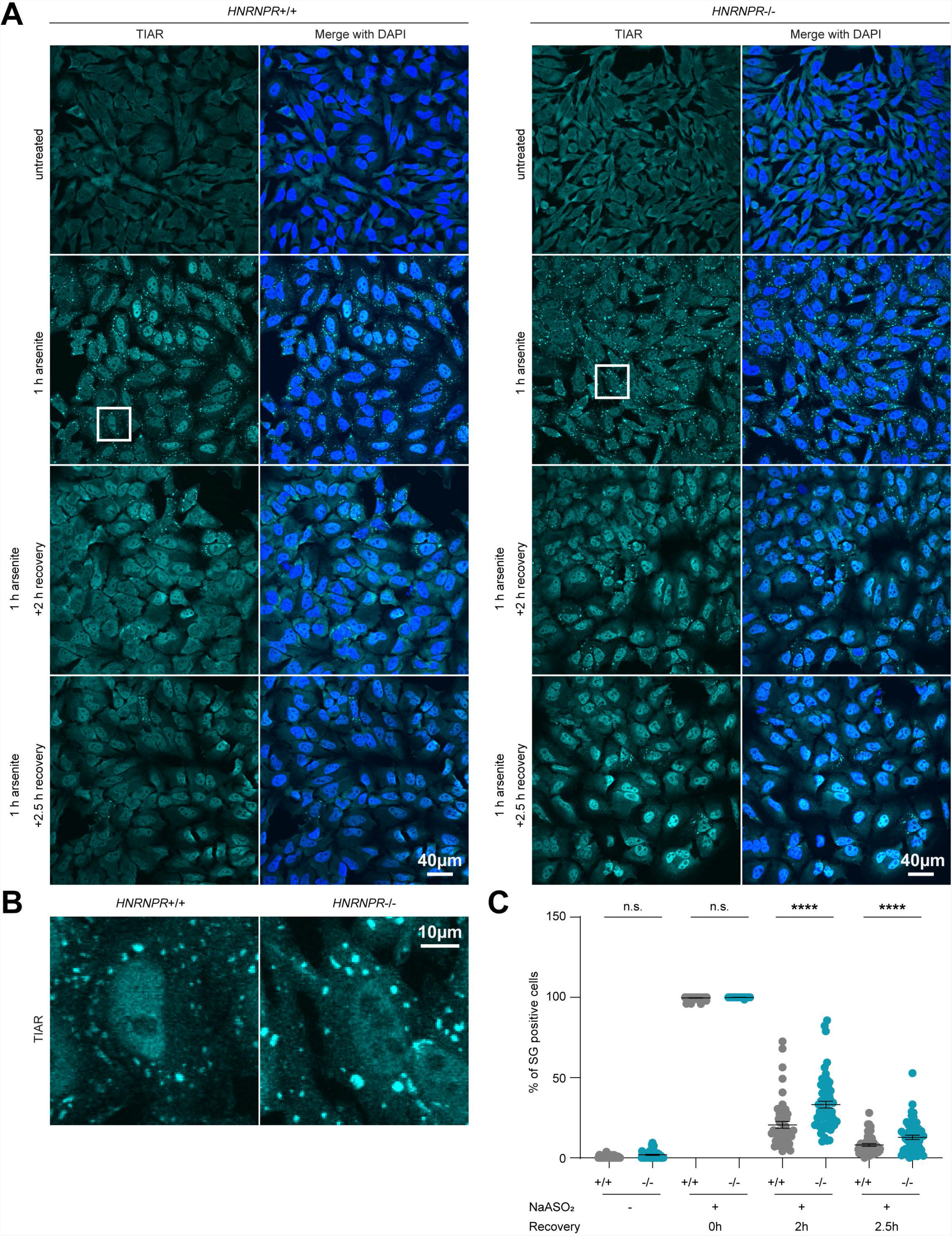
depleting hnRNP R impaires stress granule resolving. A *HNRNPR* +/+ and -/- cells were treated with water or 0.5mM sodium arsenite for one hour, meanwhile wash off sodium arsenite by adding fresh medium, Immunostaining with antibodies against TIAR to check percent of stress granule positive cells. Scale bar: 40 μm. B Zoom in the single cell of *HNRNPR* +/+ and -/- cells were treated with 0.5mM sodium arsenite for one hour. Scale bar: 10 μm. C Quantification of the percent of stress granule positive cells in Figure 6A.

**Figure 7.**
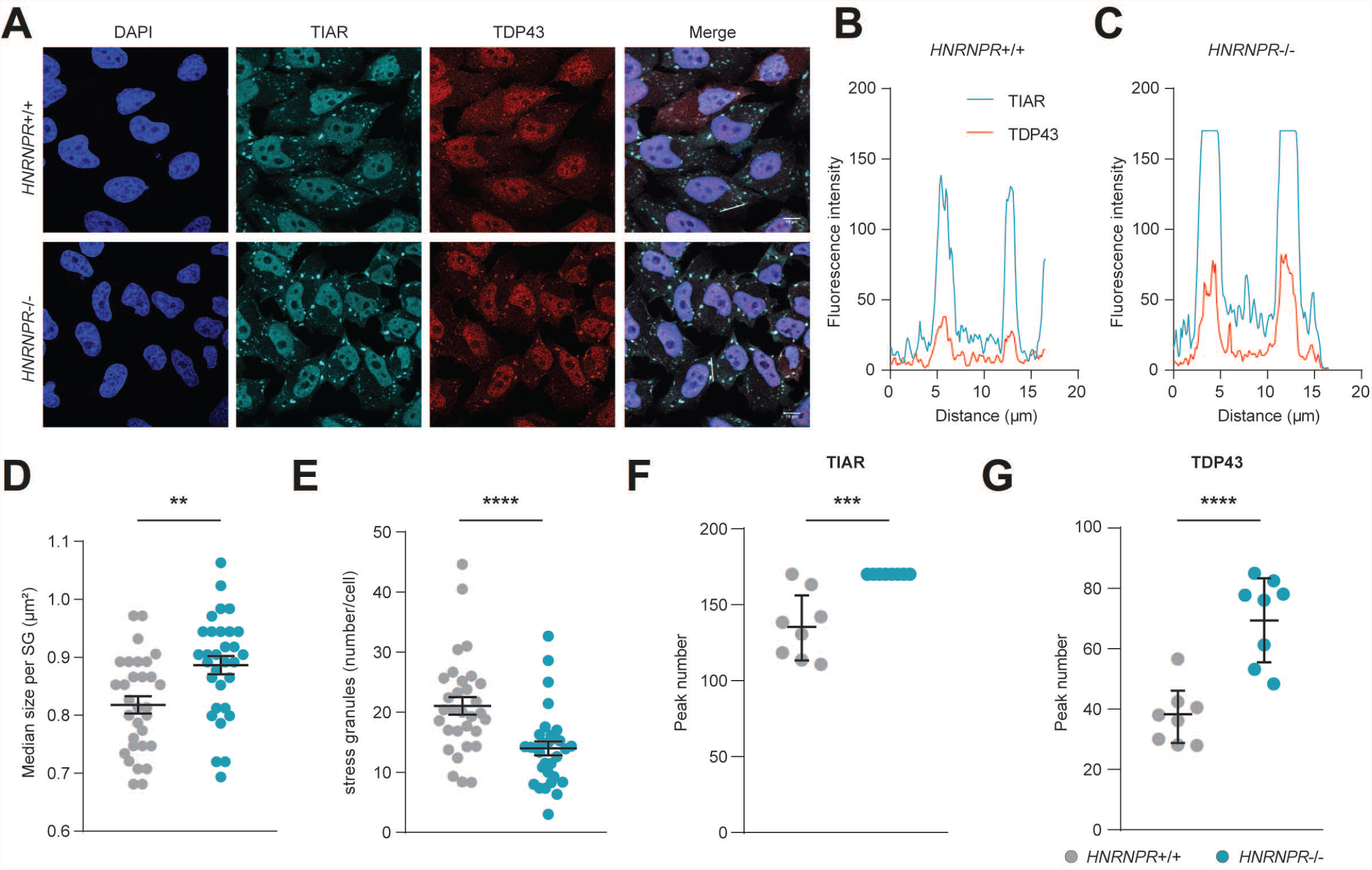
depleting hnRNP R altered TDP43 stress granule formation. A Immunostaining with antibodies against TIAR and TDP43 in *HNRNPR* +/+ and -/- cells with sodium arsenite treatment. Scale bar: 10 μm. B, C TIAR and TDP43 granule fluorescence intensity curve in *HNRNPR* +/+ and -/- cells D Quantification of the median size per stress granules in *HNRNPR* +/+ and -/- cells. Data are mean with standard deviation (SD); \*\**P* ≤ 0.01; unpaired two-tailed *t*-test (n = 3 biological replicates). E Quantification of the number of stress granules in per in *HNRNPR* +/+ and -/- cells. Data are mean with standard deviation (SD); \*\*\*\**P* ≤ 0.0001; unpaired two-tailed *t*-test (n = 3 biological replicates). F, G Quantification of the TIAR and TDP43 granule fluorescence peak number in *HNRNPR* +/+ and -/- cells. Data are mean with standard deviation (SD); \*\*\**P* ≤ 0.001; \*\*\*\**P* ≤ 0.0001; unpaired two-tailed *t*-test (n = 3 biological replicates).

### 6. depleting hnRNP R impaires stress granule resolving

Next question we want to ask whether depletion *HNRNPR* can affect stress granule recovery, base on this question, we treat *HNRNPR* +/+ and -/-cells with sodium arsenite meanwhile remove it by change with fresh medium and check the stress granule positive cells. Very interestingly, we found in *HNRNPR* -/-cells, the stress granules much difficult to resolve (Figure 6). One explain would be depletion *HNRNPR* changed stress granule physical properties, which means the internal structure and the competent of stress granule was changed.

### 7. depleting hnRNP R altered TDP43 stress granule formation

TDP43, one of the most important RNA binding proteins which mutations are associated with ALS,FTD, Parkinson disease [50] [51] [52]. Stress granules (SGs) facilitate TAR DNA-binding protein 43 (TDP-43) cytoplasmic mis localization and aggregation [53]. Stress granules are dense aggregations in the cytosol composed of proteins & RNAs that appear when the cell is under stress [54], finding out which protein or RNA regulate TDP43 recruited into stress granules still is a huge challenge. However, based on our findings, hnRNP R can regulate stress granules formation, depleting hnRNP R by knocking down or knock out can increase stress granule size meanwhile reduce stress granules number as figure 7 shows, regarding TDP43 is one competent of stress granules, which is more conducive to form aggregates after hnRNP R was abolished.

### 8. depleting hnRNP R impairs TDP43 association with ribosome

hnRNP R is one member of hnRNP family, hnRNP proteins associated with ribosomes has been wildly investigated [55] [56] (figure 8), and of course, hnRNP R is not an exception [57], although TDP43 have DNA binding activity besides RNA binding ability, but TDP43 interactome proved that TDP43 extensively associated with hnRNP proteins and translation machinery [58], but the connection between hnRNP R and TDP43 so far still not been investigated yet, here we found that dramatically induction of TDP43 binding with ribosomes after knocking out hnRNP R(figure 8), one explain would be some mRNAs can both bind hnRNP R and TDP43, and hnRNP R have the function to guide the mRNAs were translated, TDP43 itself alone cannot finish the mission to guide the mRNAs been translated without hnRNP R. Regarding TDP43 and hnRNP R mainly localized in the nuclear, another possibility would be impaired TDP43 export to cytoplasm after knocking out hnRNP R. but this need to be proved further.

**Figure 8.**
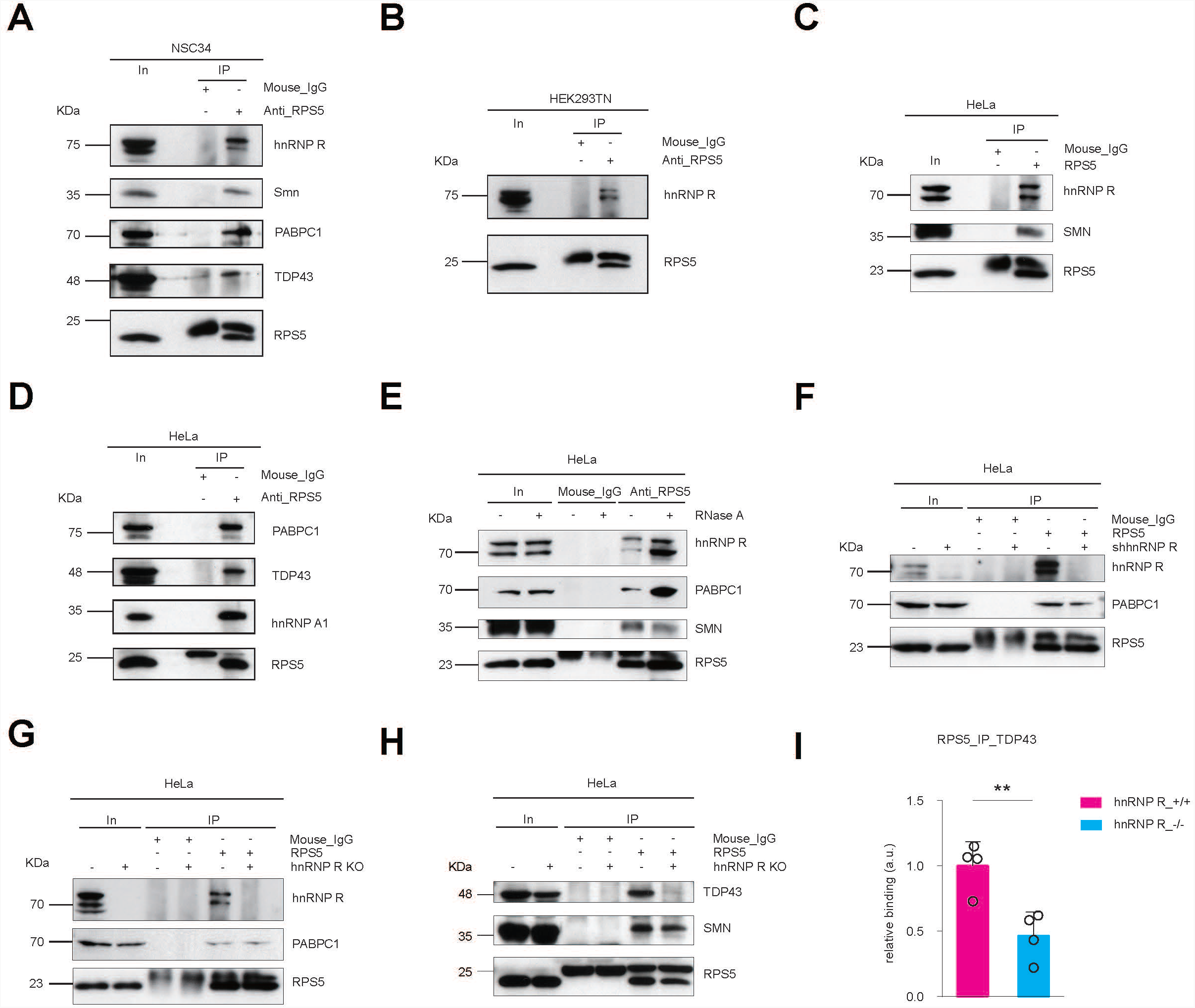
depleting hnRNP R impairs TDP43 association with ribosome. A Western blot analysis of hnRNP R, Smn, CDK9, PABPC1 and TDP43 co-immunoprecipitated by an anti-RPS5 antibody in NSC34 cells. Immunoprecipitation with mouse-IgG antibody was used as control. B Western blot analysis of hnRNP R co-immunoprecipitated by an anti-RPS5 antibody in HEK293TN cells. Immunoprecipitation with mouse-IgG antibody was used as control. C Western blot analysis of hnRNP R, SMN co-immunoprecipitated by an anti-RPS5 antibody in HeLa cells. Immunoprecipitation with mouse-IgG antibody was used as control. D Western blot analysis of PABPC1, TDP43, hnRNP A1 co-immunoprecipitated by an anti-RPS5 antibody in HeLa cells. Immunoprecipitation with mouse-IgG antibody was used as control. E Western blot analysis of hnRNP R, PABPC1, SMN co-immunoprecipitated by an anti-RPS5 antibody in HeLa cells with or without RNase A treatment. Immunoprecipitation with mouse-IgG antibody was used as control. F Western blot analysis of hnRNP R, PABPC1, co-immunoprecipitated by an anti-RPS5 antibody in control and sh*HNRNPR* HeLa cells. Immunoprecipitation with mouse-IgG antibody was used as control. G, H Western blot analysis of hnRNP R, PABPC1, TDP43, SMN co-immunoprecipitated by an anti-RPS5 antibody in *HNRNPR* +/+ and -/- cells HeLa cells. Immunoprecipitation with mouse-IgG antibody was used as control. I Quantification of TDP43 co-immunoprecipitated by an anti-RPS5 antibody in *HNRNPR* +/+ and -/- cells HeLa cells.

## DISCUSSION

This study has shown for the first time that hnRNP R regulates the stress granule formation, phenotype, and facilities the TDP43 granule formation. Western analysis revealed that TDP43 and TIAR expression was not disturbed after hnRNP R depletion. Some different mechanisms maybe involved in the stress granule formation, such as disturbed translocation from nuclear to the cytosol and vice versa. Some biochemical fractionation methods can be used to answer this question. Another possibility would because of a broader dysregulation of DNA/RNA binding proteins after depletion hnRNP R. Some proteome and transcriptome analysis methods, such as APEX can be used to Investigate the component of proteins and RNA in stress granules after hnRNP R depletion [59].

Stress granules are composed of RNA and RNA binding proteins, such as FUS, TIA1, TDP43 etc[60]. A significant percentage of motoneuron degenerative disease are caused by mutations in RNA binding proteins. Meanwhile stress granule has been found in those disease, such as TIA1 mutation ALS[61], FUS mutation ALS[62], TDP43 mutation ALS[63], c9orf72 ALS[64], additionally, stress granules have also been found in Huntington disease brain[12]. Some studies suggested that stress granules are the reason instead of the result of ALS and FTD pathogenesis, they suggested that the formation of stress granule disturbed the RNA and protein homeostasis in motoneurons. Some studies think that stress granules are the result instead of the reason, they think that the mutated RNA binding proteins disturbed the RNA splicing, RNA binding, RNA transport, RNA translation etc, In the end stress granule as protective reaction when motoneuron retaining those metabolisms defects[65].

hnRNP R is highly expressed in the brain[66], hnRNP R mutation also connected with neuron disease[67]. hnRNP R has similar structure comparing other hnRNP proteins, such as FUS, TDP43, all has RNA recognition motif (RRM), Nuclear localization signals (NLS) and prion-related domains rich in glutamine (Q) and asparagine (N) residues, although hnRNP R does not directly associated with ALS or FTD, but study found that hnRNP R can also form aggregates in FTLD-FUS brain[68]. This means that hnRNP R as a co-factor can involve in FTD. Another thing needs to mention in the study is we found depleting hnRNP R can reduce TDP43 binding with ribosomes. One explain would be without hnRNP R, TDP43 fails to guide mRNA to the ribosomes. In summary, in this study, we showed that hnRNP R can involve in stress granule regulation. potential therapy, such as overexpress hnRNP R in ALS motoneurons would be one solution to ALS treatment.

## Materials and Methods

### NSC43, HEK293TN and HeLa cell culture

NSC-34 cells (Cedarlane, cat. no. CLU140), HEK293TN cells (System Biosciences, cat. no. LV900A-1) and HeLa cells (Leibniz Institute DSMZ-German Collection of Microorganisms and Cell Cultures GmbH, DSMZ no. ACC 57) were cultured at 37°C and 5% CO_2_ in high glucose Dulbecco’s Modified Eagle Medium (DMEM; Gibco) supplemented with 10% fetal calf serum (Linaris), 2 mM GlutaMAX (Gibco) and 1% Penicillin-Streptomycin (Gibco). Cells were passaged when they were 80-90% confluent.

### Immunofluorescence staining of cultured HEK293TN and HeLa cell

For Immunofluorescence staining, 2×105 cells were grown in 10 mm Cover slips coated Greiner CELLSTAR® dish (diam. x H 35 mm x 10 mm, vented with 4 inner rings) for 2 d. For NaASO2 treatment, the medium was replaced with fresh medium containing 0.5 µM NaASO2 or H2O as control and incubated 1 hour to harvesting for staining. Cells were washed three times with prewarmed phosphate-buffered saline (PBS) and fixed with 4% paraformaldehyde (PFA) for 20 min at room temperature (RT), then washed three times with prewarmed PBS. For permeabilization, 0.3% Triton X-100 was applied for 20 min at RT followed by three washes with prewarmed PBS. Motoneurons were treated with 10% horse serum, 2% BSA in Tris-buffered saline with Tween 20 (TBS-T) for 0.5 h at RT to reduce unspecific binding followed by primary antibody incubation overnight at 4°C. The cells were washed three times with TBS-T and incubated with appropriate fluorescently labeled secondary antibodies in TBS-T for 1 h at RT. Motoneurons were washed three times with TBS-T at RT, then washed once with water. Motoneurons were embedded with Aqua-Poly/Mount (Polysciences, 18606-20). The following primary and secondary antibodies were used for immunostaining: polyclonal chicken anti-GFP (Ab13970, Abcam; 1:1,000), Mouse monoclonal TIAR antibody (610352, BD Biosciences; 1:250), Rabbit Polyclonal anti-TDP43 (C-terminal) (12892-1-AP, Proteintech; 1:250), donkey anti-chicken IgG (H+L) (Alexa 488; 703-545-155, Jackson Immunoresearch; 1:800), donkey anti-mouse IgG (H+L) (Cy 3; 715-165-151, Jackson Immunoresearch; 1:800), and donkey anti-rabbit IgG (H+L) (Alexa Fluor® 488; 711-545-152, Jackson Immunoresearch; 1:800)

### Confocal microscopy

Coronal brain slices were analyzed with an Olympus FluoView 1000 confocal laser microscope equipped with the following objectives: 40× (oil differential interference contrast, NA: 1.30) for counting the stress granules positive cells, or 60× (oil differential interference contrast, NA: 1.35) for counting stress granules numbers. Images were obtained with the corresponding Olympus FV10-ASW (RRID:<u>SCR_014215</u>) imaging software for visualization and image acquisition in a single-channel scan mode as *z* stacks, using 405, 473, 559, and 633 nm lasers. The resulting images (Olympus. oib format) were processed using ImageJ (RRID:<u>SCR_003070</u>) and projected as either maximum or average intensity (indicated in the figure legends for all images shown in this study). Finally, the data were transferred into tif format, arranged with Adobe Illustrator software (RRID:<u>SCR_010279</u>), and saved as 300 dpi png and tif files.

### Preparation of lentiviral knockdown constructs

shRNAs targeting human *HNRNPR* were cloned into a modified version of pSIH-H1 shRNA vector (System Biosciences) containing EGFP according to the manufacturer’s instructions. The following antisense sequences were used for designing shRNA oligonucleotides targeting *HNRNPR*: 5’-GTTCTGCTTCCTTGAATATGA -3’, (Supplementary Table 1). Empty pSIH-H1 expressing EGFP was used as control. Lentiviral particles were packaged in HEK293T cells with pCMV-pRRE, pCMV-pRSV, and pCMV-pMD2G as described before^48^. To assess knockdown efficiency, RNA was extracted from HeLa cells using the Trizol (Ambion) and reverse-transcribed with random hexamers using the First Strand cDNA Synthesis Kit (Thermo Fisher Scientific). Reverse transcription reactions were diluted 1:5 in water and cDNA levels measured by qPCR. Relative expression was calculated using the ΔΔCt method.

### Generation of hnRNP R knockout HeLa cell lines by prime editing

Knockout HeLa cells were generated by prime editing [49]. The sgRNA sequence 5’-CAAGGTGCAAGAGTCCACAA-3’ was identified in exon 4 of *HNRNPR* using the Broad Institute sgRNA design tool (https://portals.broadinstitute.org/gpp/public/analysis-tools/sgrna-design) and a pegRNA was designed for insertion of an in-frame stop codon. The pegRNAs were cloned into pU6-pegRNA-GG-acceptor (Addgene Plasmid #132777) [49] as following. 1 µg pU6-pegRNA-GG-acceptor was digested with Fast Digest Eco31I (IIs class) (Thermo Fisher Scientific) in a 20 µl reaction. Following agarose gel electrophoresis, the vector backbone was gel-purified using the NucleoSpin Gel and PCR Clean‐up kit (Macherey-Nagel). Oligonucleotides pegRNA3-1 and pegRNA3-2 (Appendix Tables S1 and S2) were annealed and extended in a 30 µl PCR reaction containing 0.4 µM each of pegRNA3-1 and pegRNA3-2, 3 µl of 10× Extra Buffer, 0.33 mM of each dNTP and 1 U of Taq DNA polymerase (VWR) using one cycle of 94°C for 5 min 30 s, 60°C for 30 s and 72°C for 30 s. The pegRNA was assembled and inserted into pU6-pegRNA-GG-acceptor in a 10 µl reaction containing 25 ng digested pU6-pegRNA-GG-acceptor, 1.125 µl PCR reaction from the previous step, 45 nM pegRNA3-3AGTGA (Appendix Table S1) and 5 µl NEBuilder HiFi DNA Assembly Master Mix (NEB) and incubated at 37°C for 1 h.

For transfection, 10^5^ HeLa cells were seeded in a 12-well plate in a volume of 0.5 ml of DMEM 24 h prior transfection. After 24 h, cells were co-transfected with 750 ng of pCMV-PE2 Plasmid (Addgene Plasmid #132775) and 250 ng of assembled pU6-pegRNA-GG-acceptor using Lipofectamine 2000 (Invitrogen). Cells were harvested 72 h after transfection by trypsinization using TrypLE™ Express Enzyme (Gibco), counted and diluted count to 9 cells per ml DMEM medium. 100 µl of cell suspension were added per well of a 96-well plate. Cells were grown for 7-10 d to allow formation of colonies from single cells. Colonies were trypsinized, transferred to a 48-well plate and grown for another 7 d for cell expansion. Then, cells in each well were trypsinized and split into two wells of a 24-well plate. One of these wells was used for genotyping with primers listed in Appendix Tables S1 and S2.

### hnRNP R overexpression

An expression plasmid for hnRNP R was generated as following. Total RNA was extracted from HEK293TN cells with the NucleoSpin RNA kit (Macherey-Nagel) and 1 μg RNA was reverse-transcribed with oligo-dT18 using the First Strand cDNA Synthesis Kit (Thermo Fisher Scientific). The coding sequence of full-length hnRNP R was then PCR-amplified from the cDNA using the KAPA HiFi HotStart ReadyMix (Roche) and sub-cloned into pJET1.2 using the CloneJET PCR Cloning Kit (Thermo Fisher Scientific). Using primers listed in Appendix Table S4, the hnRNP R coding sequence with overhangs was amplified from pJET1.2-hnRNP R, and the GFP coding sequence with overhangs was amplified from a pcDNA3-GFP expression plasmid. PCR products were subjected to DpnI (Thermo Fisher Scientific) digestion and, using the NEBuilder HiFi DNA Assembly Master Mix (NEB), were inserted into pcDNA3 digested with BamHI (Thermo Fisher Scientific) to generate pcDNA3-hnRNP R-GFP. For transfection, plasmids were produced using the NucleoBond Xtra Midi kit(Macherey-Nagel) and transfected using Lipofectamine 2000 (Thermo Fisher Scientific) as following. 2.5×106 HeLa cells were seeded in a T75 flask and after 24 hours transfected with 20 μg pcDNA3-hnRNP R-GFP or 15.26 μg pcDNA3-GFP. Total protein was extracted 48 h after transfection for Western blot analysis using antibodies listed in Table S5.

### Co-immunoprecipitation by using GFP_nano tap beads

24 hours prior transfection, 0.9 × 10^6^ HeLa cells were seeded in a 60 mm dish in a volume of 4 ml of DMEM 1X-Glutamax. After 24 hours, the transfection mix was set up with 5 µg of plasmid DNA,10 µL of lipofectamine 2000 and 1ml of Opti-MEM reduced Serum Medium. After 20 minutes of incubation at room temperature, each reaction was added to the seeded cells and distributed by gently pipetting up and down without getting the cells detached. Cells were harvested after 48 hours, Cells were washed once with ice-cold DPBS and collected by scraping. Cells were lysed in 400 µL lysis buffer B [10 mM HEPES (pH 7.0), 100 mM KCl, 5 mM MgCl_2_, 0.5% NP-40] on ice for 15 min. Lysates were centrifuged at 20,000×g for 15 min at 4°C. measure the protein concentration by BCA kit (thermo) and take 15ug as input for western, take 50ul as input for qPCR, 15 µl GFP-Trap Magnetic Agarose (ChromoTek GmbH) (Invitrogen) wash once with ice cold lysis buffer B, then add 300 µl lysate were added to the GFP-Trap Magnetic Agarose beads and rotated for 2 h at 4°C. Beads were washed twice with lysis buffer B and proteins were eluted in 60ul water, take 20ul for RNA extraction, the left was dissolved into 1× Laemmli buffer [50 mM Tris-HCl (pH 6.8), 1% sodium dodecyl sulfate, 6% glycerol, 1% β-mercaptoethanol, 0.004% bromophenol blue]. Proteins were size-separated by SDS-PAGE and analyzed by immunoblotting. 500ul Trizol (Ambion) were used for input and elute RNA extraction and elute RNA in the same volume of water 20ul, then take 10ul for RevertAid First Strand cDNA Synthesis Kit (Thermo Scientific), then samples 1 in 5 dilution, take 2 µL for reactions with Luminaris HiGreen qPCR Master Mix (Thermo Fisher Scientific) on a LightCycler® 96 (Roche). Relative expression was calculated using the ΔΔCt method.

### Co-immunoprecipitation by using Ribosome S5 antibody

Cells were washed once with ice-cold DPBS and collected by scraping. Cells were lysed in 1 ml lysis buffer B [10 mM HEPES (pH 7.0), 100 mM KCl, 5 mM MgCl_2_, 0.5% NP-40] on ice for 15 min. Lysates were centrifuged at 20,000×g for 15 min at 4°C. 10 µl Dynabeads Protein G and 1 µg RPS5 antibody (Santa Cruz, sc-390935) or mouse IgG control were added to 200 µl lysis buffer and rotated for 30-40 min at RT. Then 200 µl lysate were added to the antibody-bound beads and rotated for 2 h at 4°C. Beads were washed twice with lysis buffer B and proteins were eluted in 1× Laemmli buffer [50 mM Tris-HCl (pH 6.8), 1% sodium dodecyl sulfate, 6% glycerol, 1% β-mercaptoethanol, 0.004% bromophenol blue]. Proteins were size-separated by SDS-PAGE and analyzed by immunoblotting. For RNase treatment, 3 µl RNase A (Thermo Fisher Scientific) were added to 200 µl lysate and incubated at 37°C for 10 min before proceeding with co-immunoprecipitation.

### Poly(A) FISH combine with staining

HeLa cells were grown on 10 mm coverslips for 2 d, washed once with PBS and fixed with 4% PFA for 10 min. Following removal of PFA, cells were incubated with 100% cold methanol for 10 min and then with 70% ethanol for 10 min. Ethanol was aspirated and cells were incubated with 1 M Tris pH 8.0 (Thermo Scientific) for 5 min. While cells were incubated in Tris, oligo-dT probe [cy3-T(+T) TT(+T) TT(+T) TT(+T) TT(+T) TT(+T) TT(+T), where (+T) is LNA] was diluted to a final concentration of 2 ng/µl in hybridization buffer [2× SSC (Thermo Fisher Scientific), 0.1 mg/ml yeast tRNA (Sigma-Aldrich), 0.005% Ultrapure™ BSA (50 mg/mL) (Thermo Scientific), 10% dextran sulfate (Sigma-Aldrich) and 25% formamide (Sigma-Aldrich)]. Tris was aspirated and cells were incubated with diluted probe at 37°C for 5 h. After hybridization, cells were washed once with 4×SSC, once with 2×SSC, primary antibody g3bp1(1:500, Proteintech,13057-2-AP), TIAR (1:500, BD,610352) in 2xSSC + 0.01% triton-X-100 and incubate for overnight at 4C. Wash three times with 2x SSC, then incubate with secondary antibody Donkey-anti-rabbit IgG affi-pure (H+L) Jackson 711-485-152(1:1000), Donkey-anti-mouse IgG Jackson 715-175-150(1:1000) for 1hour at room temperature. and incubated with DAPI (Sigma-Aldrich) diluted 1:1,000 in Dulbecco’s phosphate-buffered saline (DPBS; Sigma-Aldrich) for 10 min. Following DAPI staining, cells were washed twice with water and coverslips were mounted.

### qPCR

Unless stated otherwise, RNA was extracted from HeLa cells using TRIzol (Thermo Fisher Scientific) and reverse-transcribed with random hexamers using the First Strand cDNA Synthesis Kit (Thermo Fisher Scientific). Reverse transcription reactions were diluted 1:5 in water and qPCR reactions were set up with Luminaris HiGreen qPCR Master Mix (Thermo Fisher Scientific) on a LightCycler® 96 (Roche). Relative expression was calculated using the ΔΔCt method. Primers are listed in Appendix Table S6.

## Stress granule quantification

The Stress granules size and number was quantified using cell profiler(https://cellprofiler.org/).

## Antibodies

Antibodies used throughout the study are listed in Supplementary Table 4.

## Statistics and reproducibility

Statistical analyses were performed using GraphPad Prism version 9 for Windows (GraphPad Software, San Diego, California USA). All results shown are representative of at least three independent experiments.

## Supporting information

primer and antibody list

## Expanded View Figure legends

**Figure EV1.**
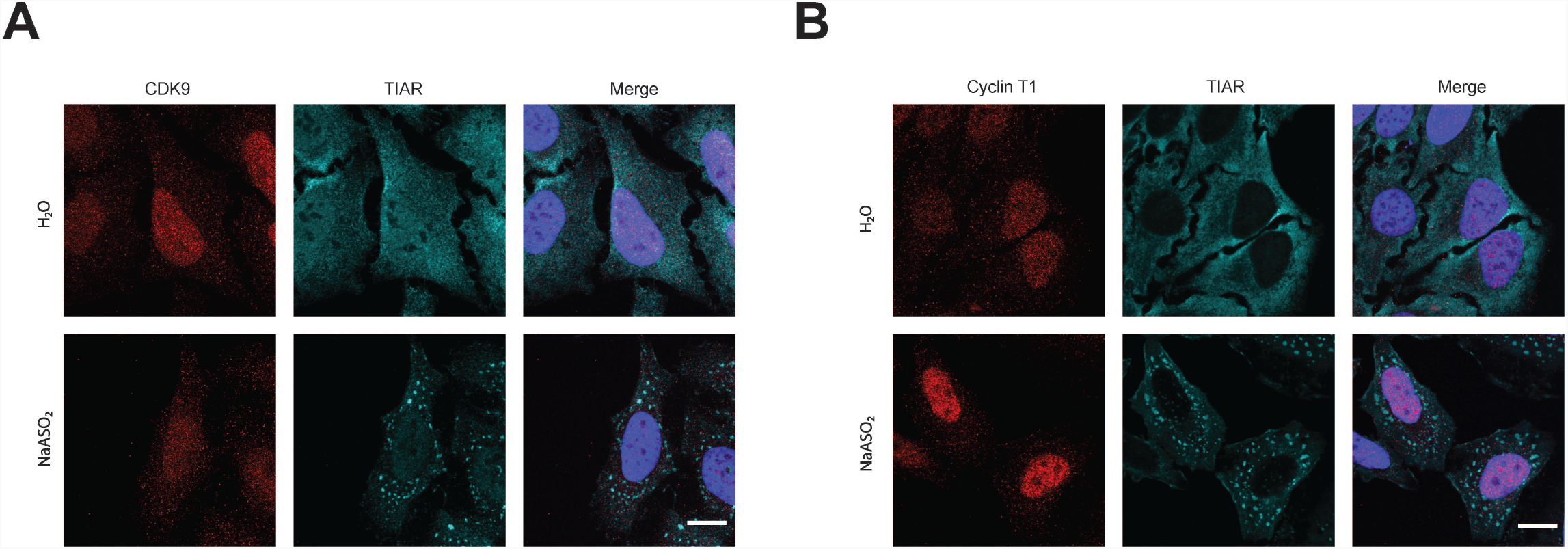
CDK9 and Cyclin T1 did not localize in stress granule. A Immunostaining with antibodies against CDK9 and TIAR in water and sodium arsenite treatment HeLa cells. Scale bar: 10 μm. B Immunostaining with antibodies against Cyclin T1 and TIAR in water and sodium arsenite treatment HeLa cells. Scale bar: 10 μm.

**Figure EV2.**
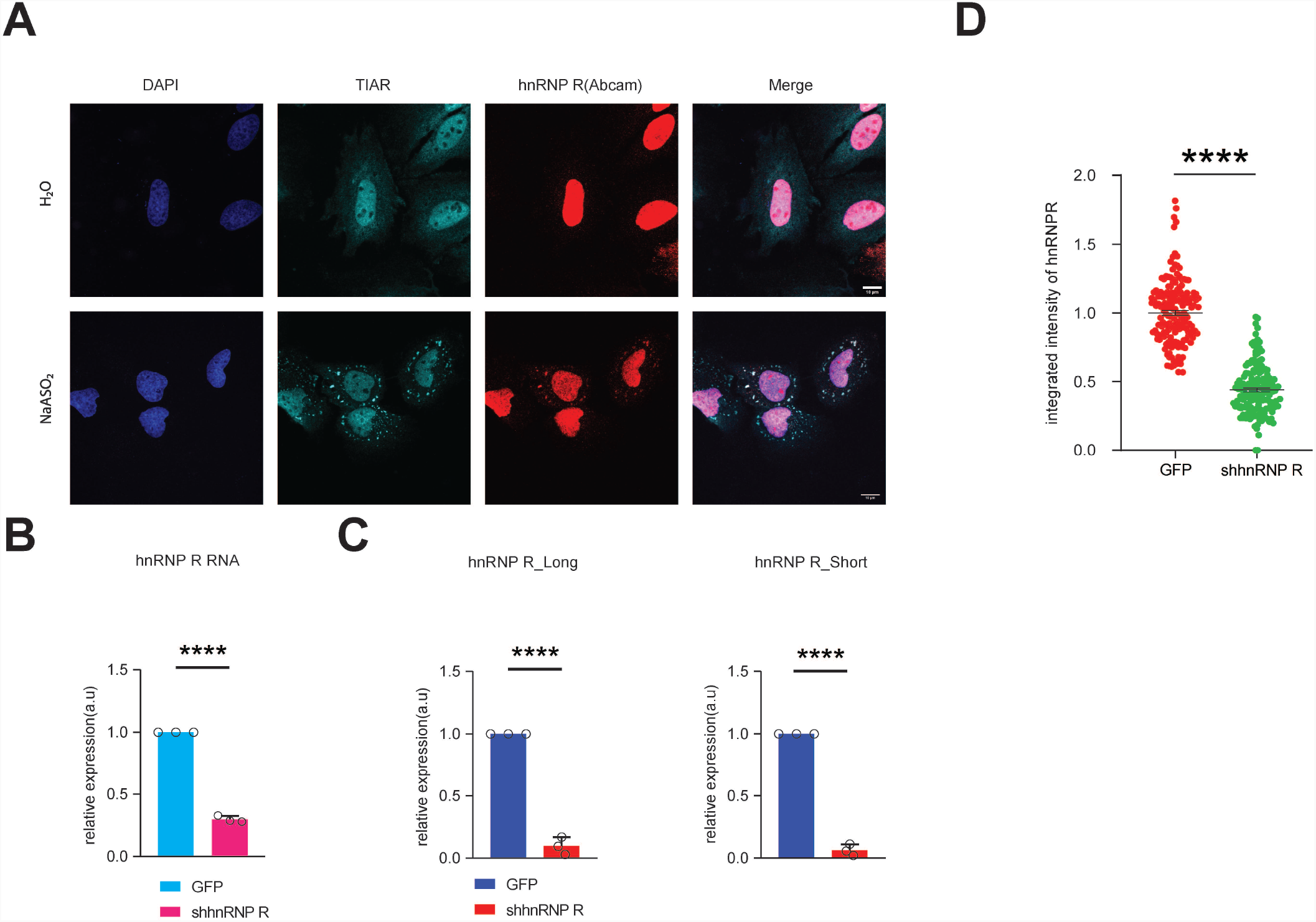
hnRNP R localized in stress granule by using different antibody. A Immunostaining with antibodies against TIAR and hnRNP R(Abcam) in water and sodium arsenite treatment HeLa cells. Scale bar: 10 μm. B qPCR analysis of *HNRNPR* RNA in control and sh*HNRNPR* HeLa cells. C Western blot analysis of hnRNP R short and long isoform protein expression in control and sh*HNRNPR* HeLa cells. D Quantification of mean fluorescence intensity of the hnRNP R signal in Figure 4(A). Data are mean with SD; \*\*\*\**P* ≤ 0.0001; Mann-Whitney test.

**Figure EV3.**
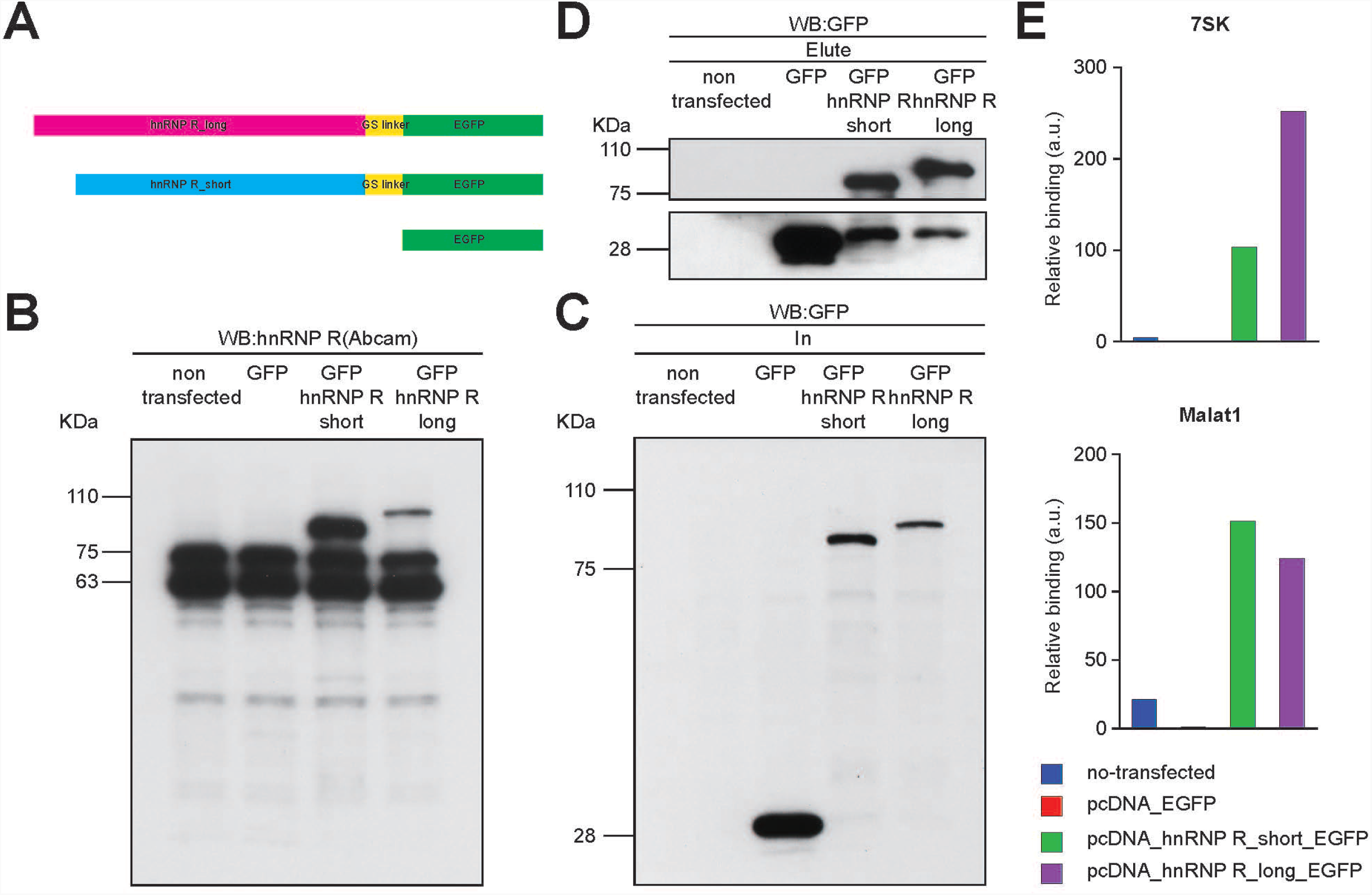
hnRNP R binds 7SK and Malat1. A Schematic of the strategy for the generation of hnRNP R_long_GFP, hnRNP R_short_GFP plasmids. B, C Western blot analysis of hnRNP R, hnRNP R_short_GFP, hnRNP R_long_GFP and GFP in HeLa cells no transfected or transfected with pcDNA3-GFP, or pcDNA3_short_GFP or pcDNA3_long_GFP. D Western blot analysis of GFP co-immunoprecipitated by GFP_nano tap beads in HeLa cells no transfected or transfected with pcDNA3-GFP, or pcDNA3_short_GFP or pcDNA3_long_GFP. E qPCR analysis of 7SK and Malat1 co-immunoprecipitated by GFP_nano tap beads in HeLa cells no transfected or transfected with pcDNA3-GFP, or pcDNA3_short_GFP or pcDNA3_long_GFP.

**Figure EV4.**
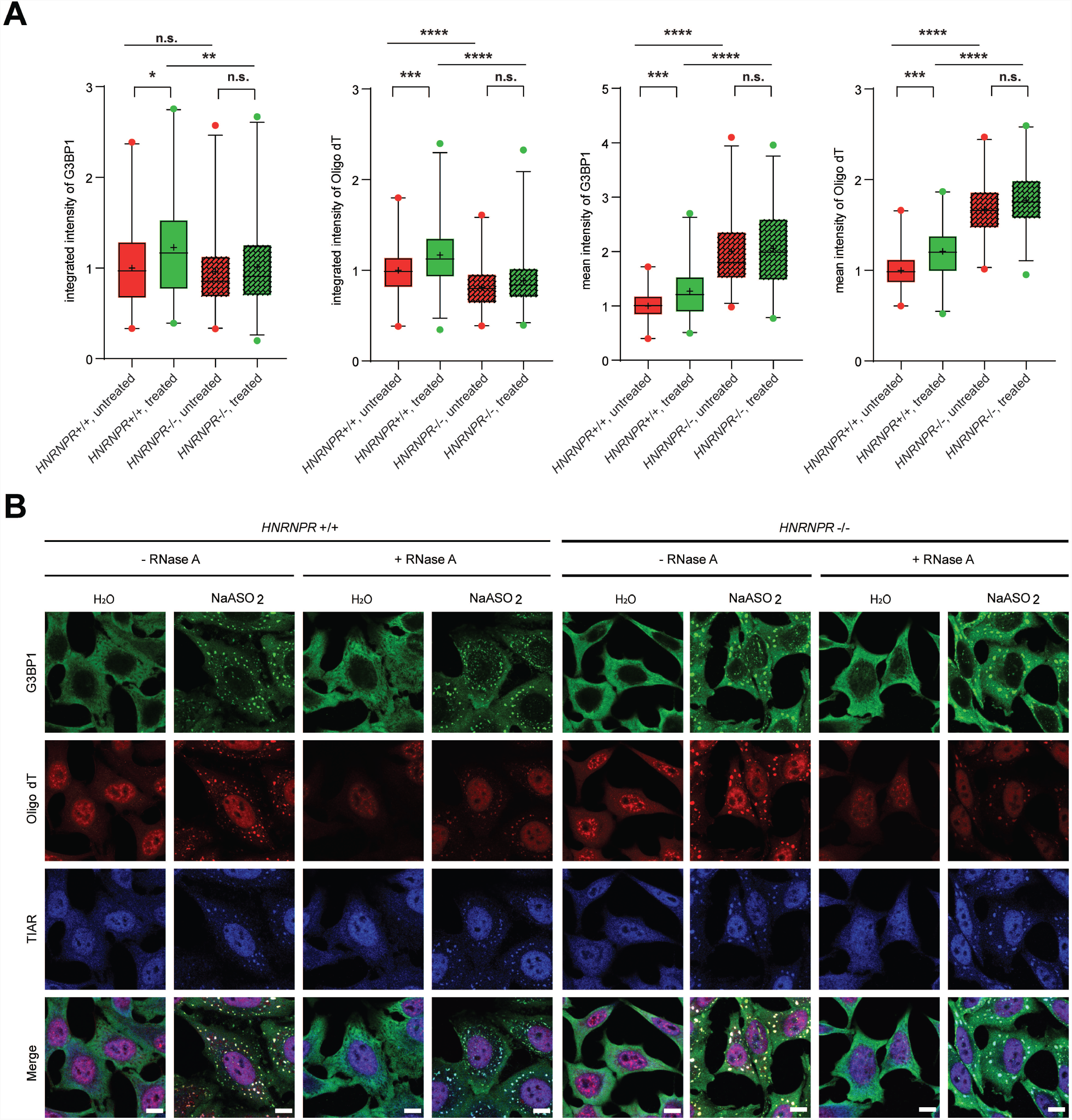
depleting hnRNP R facilitates RNA granule formation. A Quantification of the integrated and mean intensity of G3BP1 and Oligo dT in per in *HNRNPR* +/+ and -/- cells with or without RNase A treatment. Data are mean with standard deviation (SD); n.s. P > 0.05; *P ≤ 0.05; **P ≤ 0.01; ***P ≤ 0.001; \*\*\*\**P* ≤ 0.0001; unpaired two-tailed *t*-test (n = 3 biological replicates). B Poly(A) FISH and Immunostaining with antibodies against TIAR and G3BP1 in *HNRNPR* +/+ and -/- cells with or without sodium arsenite and RNase A treatment. Scale bar: 10 μm.

