## Supplementary material for "hnRNP R is a regulator of stress granule formation": primer and antibody list

Changhe Ji

**Table of Contents:**

Appendix Table S1 - Sequences of oligonucleotides for shRNA cloning

Appendix Table S2 - Sequences of oligonucleotides for prime editing

Appendix Table S3 - Sequences of oligonucleotides for qPCR

Appendix Table S4 - Sequences of oligonucleotides for *HNRNPR* overexpression

Appendix Table S5 - Antibodies

### SUPPLEMENTARY TABLES

#### Supplementary Table 1: Sequences of oligonucleotides for shRNA cloning

| Name | Sequence (5'-3') |
|---|---|
| Sh*HNRNPR*_F | GAT CCG TTC TGC TTC CTT GAA TAT GAT CAA GAG TCA TAT TCA AGG AAG CAG AAC TTT TTG |
| Sh*HNRNPR*_R | AAT TCA AAA AGT TCT GCT TCC TTG AAT ATG ACT CTT GAT CAT ATT CAA GGA AGC AGA ACG |

#### Supplementary Table 2: Sequences of oligonucleotides for prime editing

| Name | Sequence (5'-3') |
|---|---|
| pegRNA3-1 | TAT CTT GTG GAA AGG ACG AAA CAC CGC AAG GTG CAA GAG TCC ACA AGT TTT AGA GCT AGA AAT AGC |
| pegRNA3-2 | GCA CCG ACT CGG TGC CAC TTT TTC AAG TTG ATA ACG GAC TAG CCT TAT TTT AAC TTG CTA TTT CTA GCT CTA AAA C |
| PegRNA3-3AGTGA | TGA AAA AGT GGC ACC GAG TCG GTG CTC AGG TCC CTT TGT CAC TTG GAC TCT TGC ACC TTT TTT TAA GCT TGG GCC GCT CGA G |
| EXON4_F | TCCGACATCTGGCAAAGACA |
| EXON4_F (for AGTGA) | AGG TGC AAG AGT CCA AGT GA |
| EXON4_R | GGTCAAATGCCTCTTTCCATGT |

#### Supplementary Table 3: Sequences of oligonucleotides for qPCR

| Gene | Forward primer (5'–3') | Reverse primer (5'–3') |
|---|---|---|
| 7SK | ATTGATCGCCAGGGTTGATTCG | ATGGACCTTGAGAGCTTGTTTG |
| Malat1 | TGCAGTGTGCCAATGTTTCG | AGTCTGCTGTTTCCTGCTCC |
| hnRNP R | AAG TCA CAG AGG GTT TGG TG | GCT TGT GCT GCT GAC TTG TG |

#### Supplementary Table 4: Sequences of oligonucleotides for *HNRNPR* overexpression

| Name | Sequence (5'-3') |
|---|---|
| *HNRNPR*_Long_F | CCA AGC TTG GTA CCG AGC TCG GAT CCG TCG CCA CCA TGG CTA ATCAGG TGA ATG G |
| *HNRNPR*_R | CGC TAC CGC TAC CGC TGC CGC TAC CCT TCC ACT GTT GCC CAT AAG |
| EGFP_F | CGG CAG CGG TAG CGG TAG CGG CAG CGT GAG CAA GGG CGA GGA GCT |
| EGFP_R | CAC TGG CGG CCG TTA CTA GTG GAT CCC TAC TTG TAC AGC TCG TCC A |

**Supplementary Table 5: Antibodies used for Western blotting (WB), immunoprecipitation (IP), immunofluorescence (IF) or motoneuron cell culture**

| Antibody | Application | Dilution | Source | Identifier |
|---|---|---|---|---|
| Mouse monoclonal Anti-Ribosomal Protein S5 (A-8) | WB<br>IP | 1:1000 | Santa Cruz | Cat# sc-390935;<br>RRID:AB_2713966 |
| Rabbit polyclonal anti-TDP-43 (C-terminal) | WB | 1:1000 | Proteintech | Cat# 12892-1-AP;<br>RRID: AB_2200505 |
| Mouse monoclonal anti- SMN | WB | 1:2000 | BD | Cat# 610647.<br>RRID: AB_397973 |
| Rabbit polyclonal anti-LARP7 | IF | 1:100 | Proteintech | Cat#17067-1-AP;<br>RRID: AB_2132693 |
| Rabbit polyclonal anti-MePCE | IF | 1:200 | Proteintech | Cat#14917-1-AP;<br>RRID: AB_2250635 |
| Rabbit polyclonal anti-hnRNP R | WB<br>IF | 1:4000<br>1:200 | Abcam | Cat#ab30930;<br>RRID:AB_2295532 |
| Rabbit polyclonal anti-hnRNP R Antibody (N-term) | WB | 1:1000 | Abgent | Cat#AP17239a;<br>RRID: AB_11136203 |
| Rabbit polyclonal anti-hnRNP R Antibody(C-term) | IF | 1:200 | Sigma | Cat# HPA026092;<br>RRID: AB_1850885 |
| Rabbit polyclonal anti-Cyclin T1 (D1B6G) | WB<br>IP | 1:1000 | Cell Signaling Technology | Cat# 81464S;<br>RRID:AB_2799973 |
| Mouse monoclonal anti-Cyclin T1 (clone C-6) | WB<br>IF | 1:100<br>1:100 | Santa Cruz | Cat#sc-271575;<br>RRID: AB_10650141 |
| Rabbit polyclonal anti-Cdk9 (C12F7) | WB<br>IF | 1:1000<br>1:250 | Cell Signaling Technology | Cat#2316S<br>RRID: AB_2291505 |
| Mouse monoclonal anti-Cdk9 (clone D-7) | WB<br>IP | 1:100 | Santa Cruz | Cat#sc-13130;<br>RRID:AB_627245 |
| Mouse monoclonal anti-TIAR | IF<br>WB | 1:400<br>1:2000 | BD | Cat#610352;<br>RRID:AB_397742 |
| Rabbit polyclonal anti-G3BP1 | IF | 1:500 | Proteintech | Cat# 13057-2-AP;<br>RRID: AB_2232034 |
| Mouse monoclonal anti-hnRNP A1 (clone 4B10) | WB | 1:2000 | Santa Cruz | Cat#sc-32301;<br>RRID: AB_627729 |
| Mouse monoclonal anti-G3BP1 | IF | 1:500 | BD | Cat# 611127;<br>RRID:AB_398438 |
| Rabbit polyclonal anti-HEXIM1 | IF | 1:100 | Bethyl | Cat#A303-113A;<br>RRID:AB_10892626 |
| Mouse monoclonal anti-PABP (10E10) | WB<br>IF | 1:1000<br>1:250 | Santa Cruz | Cat#sc-32318;<br>RRID:AB_628097 |

| Antibody | Application | Dilution | Source | Identifier |
|---|---|---|---|---|
| Mouse IgG control | IP | | Santa Cruz | Cat#sc-2025; RRID:AB_737182 |
| Goat polyclonal anti-Mouse,Peroxidase conjugated | WB | 1:5000 | Jackson ImmunoResearch | Cat#115-035-146. RRID: AB_2307392 |
| Donkey polyclonal anti-Rabbit, Peroxidase conjugated | WB | 1:5000 | Jackson ImmunoResearch | Cat#711-035-152; RRID:AB_10015282 |
| Donkey polyclonal anti-Goat,Peroxidase conjugated | WB | 1:5000 | Jackson ImmunoResearch | Cat#705-035-003; RRID: AB_2340390 |
| Donkey polyclonal anti-Chicken, Alexa Fluor® 488 conjugated | IF | 1:800 | Jackson ImmunoResearch | Cat#703-545-155; RRID: AB_2340375 |
| Donkey polyclonal anti-Mouse, Cy™3 conjugated | IF | 1:800 | Jackson ImmunoResearch | Cat#715-165-151. RRID: AB_2315777 |
| Donkey polyclonal anti-Rabbit, Cy™5 conjugated | IF | 1:800 | Jackson ImmunoResearch | Cat#711-175-152; RRID: AB_2340607 |
